# Multi-area single-cell calcium imaging dataset of the mouse cortex across wakefulness, sleep, and anesthesia

**DOI:** 10.64898/2026.07.20.739676

**Authors:** Ikumi Oomoto, Daiki Kiyooka, Masafumi Oizumi, Masanori Murayama

**Affiliations:** Laboratory for Haptic Perception and Cognitive Physiology, RIKEN Center for Brain Science, Wako-shi 351-0198, Saitama, Japan; Department of General Systems Studies, Graduate School of Arts and Sciences, The University of Tokyo, 3-8-1 Komaba, Meguro-ku, Tokyo 153-8902, Japan

## Abstract

We present a reusable dataset of neuronal population activity from multiple cortical areas, recorded from layers 2/3 of the mouse cortex at single-cell resolution during wakefulness, natural sleep (including NREM and REM sleep), and isoflurane anesthesia. Using wide-field two-photon microscopy, we recorded approximately 4,000 to 10,000 neurons per session at 7.65 Hz and provided the spatial coordinates of individual neurons. The repository provides both processed datasets and the corresponding raw imaging movies (TIFF) and electrophysiological recordings (MATLAB format). Processed data are distributed in MATLAB format and include Δ F/F fluorescence signals, deconvolved spike estimates, Gaussian-smoothed spike estimates, behavioral state annotations, and metadata. This dataset supports reuse in studies of cortical population dynamics, brain-state-dependent activity, and spatially distributed neuronal organization. It should also be useful for method development, benchmarking, and comparative analyses of large-scale neuronal activity across physiological and pharmacological brain states.

## Background & Summary

Understanding how cortical activity is organized across brain states such as wakefulness, sleep, and anesthesia is a central question in systems neuroscience. These states exhibit distinct patterns of neuronal activity while arising from the same underlying cortical circuitry, providing a framework for studying large-scale neural dynamics^1,2^. While local circuit properties and sensory responsiveness are partly preserved across states^3–6^, their network-level organization differs substantially. Wakefulness is characterized by desynchronized activity and sustained inter-areal coordination, whereas NREM sleep shows slow oscillations and alternating UP and DOWN states, and REM sleep exhibits partial desynchronization under distinct neuromodulatory conditions^7–9^. Anesthesia further disrupts long-range coordination and promotes spatially restricted activity patterns^10^. These state-dependent differences have been extensively characterized using macroscopic recording techniques such as functional magnetic resonance imaging (fMRI), electroencephalography (EEG), and electrocorticography (ECoG), which capture large-scale changes in brain activity and inter-areal coordination across states. Because these approaches measure neural activity indirectly or at coarse spatial resolution, they do not resolve the activity of individual neurons, the fundamental units of neural information processing. Understanding how large-scale dynamics arise from individual activity, therefore, requires measurements at single-cell resolution, ideally combined with broad spatial coverage and reliable brain-state annotation.

Wide-field two-photon microscopy, including the FASHIO-2PM platform developed in our group through a series of prior studies, has recently made this possible by combining broad spatial coverage with single-cell resolution, thereby enabling simultaneous recording from thousands of identified neurons across multiple neighboring cortical areas within a single field of view^11–14^. This capability enables examination of how state-dependent dynamics, inter-areal coordination, and spatially distributed network structure are expressed at the level of individual cells and across mesoscale cortical space. Despite these advances, wide-field two-photon microscopy has not yet been widely adopted, and consequently, publicly available resources or datasets containing large-scale single-cell activity data across multiple brain regions with brain-state annotation remain limited.

Here, we present a dataset that simultaneously delivers the three properties that prior modalities have not provided together — single-cell resolution, broad contiguous cortical coverage, and reliable brain-state annotation within the same recordings — comprising activity from thousands of layers 2/3 neurons across multiple cortical areas during wakefulness, natural sleep, and isoflurane anesthesia. Each recording session provides approximately 4,000–10,000 neurons at 7.65 Hz, together with spatial coordinates and cortical region labels for every neuron. Processed activity is distributed as three complementary representations — Δ F/F fluorescence signals, deconvolved spike estimates, and Gaussian-smoothed activity traces — alongside aligned behavioral state annotations that distinguish wakefulness, quiet wakefulness, NREM sleep, and REM sleep in sleep sessions, and wakefulness and isoflurane anesthesia in anesthesia sessions. By combining broad cortical coverage, cellular resolution, and brain-state annotation within the same recordings, this resource enables studies of cortical population dynamics across brain states, benchmarking of analytical pipelines, and comparative analyses of spatially organized neuronal activity across physiological and pharmacological conditions.

This repository comprises the complete dataset associated with our previous study^12^, including both processed recordings and the corresponding raw imaging and electrophysiological data. The processed datasets are provided in a structured format to facilitate reuse. It is intended as a resource for analyses of cortical population dynamics across brain states, for benchmarking and validation of analytical methods, and for comparative studies of spatially organized neuronal activity across physiological and pharmacological conditions.

## Methods

### Dataset overview

This Data Descriptor reports all processed recording sessions used in our previous study of cortical activity across brain states. The dataset consists of wide-field two-photon calcium imaging recordings obtained from layer 2/3 neurons of the mouse cortex during wakefulness, natural sleep, and isoflurane anesthesia, together with spatial coordinates, cortical region labels, behavioral state annotations, frame annotations, and metadata. Each shared file corresponds to one continuous recording session and contains processed fluorescence signals, deconvolved spike estimates, and Gaussian-smoothed activity traces. In addition to the processed data, the repository also provides the corresponding raw imaging movies in TIFF format and raw EEG/EMG recordings in MATLAB (.mat) format, enabling users to reproduce preprocessing and behavioral-state annotation from the original recordings.

Because the sleep and anesthesia datasets were acquired from different animal cohorts and at different circadian phases, direct quantitative comparisons between these experimental conditions should be interpreted with appropriate caution. The primary purpose of this dataset is to facilitate analyses within each recording paradigm while enabling exploratory comparisons across conditions.

### Animals

All animal experiments were performed in accordance with institutional guidelines and were approved by the RIKEN Animal Care and Use Committee (approval number: W2024-2-013(2)). Wild-type C57BL/6JJmsSlc mice (Japan SLC, Shizuoka, Japan) of both sexes were used. Mice used for sleep recordings were 12–24 weeks old, and mice used for anesthesia recordings were 28–62 weeks old. Animals were housed under a 12 h light/12 h dark cycle with ad libitum access to food and water.

All animal experiments were performed in accordance with the ARRIVE guidelines for reporting animal research.

### AAV vector preparation and neonatal transduction

The calcium indicator G-CaMP7.09^15^ was expressed using an adeno-associated virus (AAV-DJ-Syn-G-CaMP7.09-WPRE). The construct was generated by subcloning G-CaMP7.09 into a synapsin I (SynI)-expressing vector, and the virus was produced as previously described^16^. Neonatal mice (postnatal day 0–2) were anesthetized by cryoanesthesia for 2–3 min and mounted in a custom neonatal holder. AAV diluted to 4.0 × 10^12^ genome copies/mL in phosphate-buffered saline was injected into the neonatal cortex (4 μL per pup) through a glass pipette with a tip diameter of 45–50 μm. The injection site was positioned in the frontal cortex at a depth of 250–300 μm. The injection procedure was completed within 10 min after induction of cryoanesthesia, and pups were rewarmed before being returned to their home cage.

### Cranial window implantation and electrode placement

For surgery, mice were anesthetized with isoflurane (2%) and subsequently received medetomidine/midazolam/butorphanol (MMB)^17^. Body temperature was maintained at 36– 37°C during surgery using a feedback-controlled heating pad, and ophthalmic ointment was applied to prevent corneal drying. After scalp removal, a 5-mm-diameter craniotomy was made over the right hemisphere above an area including the primary somatosensory cortex, and it was covered with a 5-mm-diameter No. 1 coverslip sealed with dental cement. Cranial window implantation and electrode placement for EEG/EMG recordings were performed as previously described^18,19^. EEG screws were implanted over the lateral secondary visual area (V2L) in the right hemisphere and over the parietal cortex in the left hemisphere, with cerebellar screws used as references. For EMG recording, a flexible wire electrode was implanted into the neck muscle. A stainless-steel headplate was fixed to the skull over the cerebellum, and the exposed skull, EEG screws, and EMG wires were covered with dental cement. After surgery, mice received atipamezole hydrochloride (0.12 mg/kg body weight) and were allowed to recover on a heating pad.

### Macroscopic imaging and cortical region annotation

After surgery, skull images, including the cranial window, were acquired using one-photon macroscopic imaging to support spatial annotation of the two-photon field of view. Images were acquired with a field of view of 13.3 × 13.3 mm and a resolution of 1,024 × 1,024 pixels after 2 × spatial binning. The excitation intensity, camera gain, and exposure time were adjusted to avoid pixel saturation. Cortical region boundaries within the two-photon imaging field were estimated with reference to the Allen Mouse Common Coordinate Framework. To account for the mouse head’s tilt during imaging, the atlas was geometrically transformed to align with the imaging plane. The atlas was first coarsely aligned to the one-photon macroscopic images using anatomical landmarks, including bregma, lambda, the anterior tip of the olfactory bulb, and the boundary between the olfactory bulb and the cerebral cortex, and then refined by matching vascular patterns between the macroscopic images and the two-photon field of view. A representative example of the atlas registration is shown in Fig. 1a. Based on this registration, each ROI was assigned a cortical region label, which is provided in the shared dataset as ROIs.atlas, together with the centroid coordinates of each ROI (ROIs.Centroid). These annotations were used to relate single-neuron activity to spatially contiguous cortical areas within the imaged 3 × 3 mm field of view. Therefore, the cortical region labels provided in ROIS.atlas should be regarded as approximate reference annotations rather than definitive anatomical assignments. This uncertainty is expected to be greatest for ROIs located near cortical-area boundaries and reflects both the manual registration procedure and inter-animal anatomical variability that is not represented in the reference atlas.

**Figure 1.**
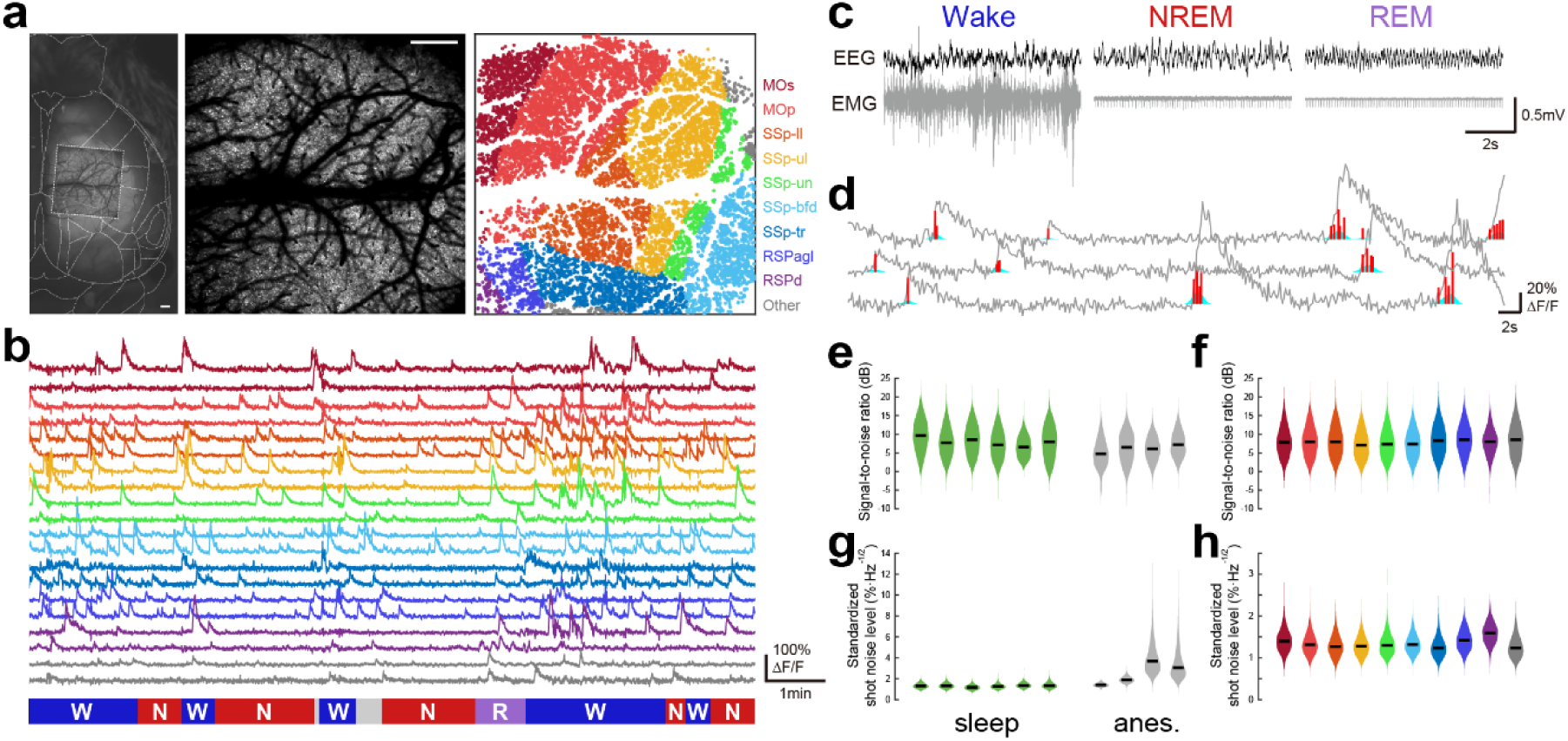
Overview of the dataset and data quality. (a) Representative atlas-registration overlay showing the approximate alignment between the macroscopic image, the Allen Mouse Common Coordinate Framework, and the two-photon imaging field (left). The two-photon field of view is outlined by a dashed square. Representative field-of-view image (center) and detected neuronal ROIs colored according to cortical region (right). Scale bars, 0.5 mm. (b) Example Δ F/F activity traces from 20 randomly selected neurons. Colors indicate the cortical region assigned to each ROI. Corresponding behavioral state annotations are shown below (W, wakefulness; N, NREM sleep; R, REM sleep; gray, quiet wakefulness). (c) Representative EEG and EMG recordings during wakefulness, NREM sleep, and REM sleep. EEG traces are low-pass filtered for display purposes only. (d) Example traces illustrating signal processing steps for three randomly selected neurons. Gray, Δ F/F fluorescence signal; red, deconvolved spike estimate; cyan, Gaussian-smoothed deconvolved signal. (e) Distribution of signal-to-noise ratio (SNR) across neurons after quality control. Each violin plot represents one recording session (n = 6 wakefulness-sleep sessions and n = 4 wakefulness-anesthesia sessions). Green indicates wakefulness-sleep sessions, and gray indicates wakefulness-anesthesia sessions. (f) Distribution of SNR across neurons after quality control, shown separately for each cortical region. Colors correspond to the cortical regions shown in (a). (g) Distribution of standardized shot noise levels across neurons after quality control. Each violin plot represents one recording session. Green indicates wakefulness-sleep sessions, and gray indicates wakefulness-anesthesia sessions. (h) Distribution of standardized shot noise levels across neurons after quality control, shown separately for each cortical region. Colors correspond to the cortical regions shown in (a). Cortical region abbreviations: MOs, secondary motor area; MOp, primary motor area; SSp-ll, primary somatosensory area, lower limb; SSp-ul, primary somatosensory area, upper limb; SSp-un, primary somatosensory area, unassigned; SSp-bfd, primary somatosensory area, barrel field; SSp-tr, primary somatosensory area, trunk; RSPagl, retrosplenial area, agranular lateral; RSPd, retrosplenial area, dorsal. Regions containing fewer than 100 neurons were grouped as “Other”.

### Wide-field two-photon calcium imaging

In vivo calcium imaging was performed using a custom-designed wide-field two-photon microscope (FASHIO-2PM)^11^. To monitor spontaneous sleep-wake behavior under the microscope, mice were head-fixed in a custom stage box and covered with an enclosure to maintain body temperature; the floor of the box was covered with home-cage bedding. Mice were gradually habituated to head fixation during the light phase, with the fixation duration increased from 1 min to 2 h per day. More than 2 weeks of habituation enabled spontaneous wakefulness, NREM sleep, and REM sleep in the recording setup. Before each imaging session, the cranial window was cleaned with acetone, and EEG/EMG electrodes were connected. Imaging was performed from layer 2/3 neuronal somata at a depth of 120–150 μm below the pial surface. G-CaMP7.09 was expressed under the synapsin promoter, and the recorded population is therefore expected to be predominantly excitatory^11,16^. G-CaMP7.09 was excited at 920 nm with a tunable Ti:sapphire laser, and fluorescence was detected with a GaAsP photomultiplier tube in the 515–565 nm range. Imaging was acquired at 7.65 Hz with 2,048 × 2,048 pixels or 1,024 × 1,024 pixels over a contiguous 3 × 3 mm field of view encompassing multiple cortical areas. Laser power at the front of the objective was typically 60–80 mW. Sleep recordings were obtained during the light phase (zeitgeber time 3–7), and anesthesia recordings were obtained during the dark phase (zeitgeber time 15–19). For anesthesia experiments, mice were recorded during wakefulness for 20–60 min and then anesthetized with 0.6% isoflurane. Raw movies were saved as 16-bit monochrome TIFF files.

### EEG/EMG recording and brain-state annotation

EEG and EMG were recorded simultaneously with two-photon imaging. Signals were amplified and filtered (EEG, 0.1–100 Hz; EMG, 5–300 Hz), digitized at 1 kHz using a 16-bit analog-to-digital converter, and acquired with Clampex software. Scan timing signals from the microscope were digitized in the same system to align EEG/EMG data with imaging frames. Behavioral state classification was performed offline from EEG and EMG signals recorded from the same hemisphere as imaging, basically as described in previous reports^18,19^. In each 4-s sliding window, the root-mean-square (RMS) of the EMG signal, EEG delta power (1–4 Hz, normalized to 1–50 Hz), and EEG theta power (6–9 Hz, normalized to 1–4 Hz) were computed. Brain states were classified as wakefulness when EMG RMS exceeded a manually defined threshold, NREM sleep when EMG RMS was below threshold, and the delta/theta ratio exceeded 0.3, and REM sleep when EMG RMS was below threshold, and the delta/theta ratio was below 0.3. Because head fixation reduced EMG variability, some immobile wake epochs were initially misclassified as REM and were manually corrected by visual inspection. Epochs shorter than 12 s were excluded and merged with adjacent states.

### Image preprocessing and ROI extraction

Imaging movies were first corrected for motion using NoRMCorre^20^. Regions of interest (ROIs) corresponding to neuronal somata were then identified using the low-computational-cost cell detection (LCCD) algorithm^21^. For each ROI, the fluorescence time course was computed as the mean fluorescence across pixels within the ROI. Neuropil-corrected somatic fluorescence was estimated as F(t) = F_ROI_(t) − *r* × F_neuropil_(t), where *r* = 0.7 following previous studies^22–24^, and F_neuropil_(t) was defined as the mean fluorescence between 4 and 9 pixels from the ROI boundary, excluding pixels belonging to any ROI.

### Fluorescence signal processing

The relative fluorescence change was calculated as Δ F/F = (F(t) − F0(t))/F_0_(t), where F_0_(t) was estimated as the 8th percentile of the fluorescence distribution within a ±30 s window around each time point^25^. The baseline was then further corrected using the estimate_baseline function implemented in Suite2P^26^. The processed dataset includes Δ F/F fluorescence signals, deconvolved spike estimates, and Gaussian-smoothed activity traces for each neuron.

### Noise metrics

For technical validation, we quantified two complementary noise-related measures for each ROI: the signal-to-noise ratio (SNR) of the Δ F/F trace and the standardized shot noise level^27^. SNR was used as part of the ROI quality-control procedure, whereas standardized shot noise levels were used to assess the consistency of data quality across sessions and cortical regions (Fig. 1d–g). Standardized shot noise levels were calculated following the definition of Rupprecht et al.^27^ and were used for descriptive comparison rather than ROI exclusion.

### Quality control, deconvolution, and data packaging

To exclude contaminated or low-quality ROIs, multiple quality-control criteria were applied. The signal-to-noise ratio (SNR) of each ROI was calculated in MATLAB, where the signal was defined as the Δ F/F component below 0.5 Hz and the noise as the component above 0.5 Hz. The 0.5 Hz threshold was selected empirically based on comparisons of power spectral densities from recordings with clear neuronal calcium activity and recordings dominated by noise, which showed that biologically meaningful calcium signals were predominantly concentrated below this frequency under our imaging conditions. ROIs were retained only if they satisfied all of the following criteria: (1) SNR within the top 90% of ROIs, (2) absence of obscuration by nearby blood vessels, (3) Δ F/F range between −1 and 10, (4) at least one calcium transient during recording, (5) neuropil fluorescence lower than somatic fluorescence, (6) fewer than 45% of frames with Δ F/F below the mean, and (7) correlation between Δ F/F and the square root of EMG below 0.2. Blood-vessel detection was performed semi-automatically in ImageJ. After quality control, Δ F/F traces were deconvolved using OASIS^28^ and deconvolved traces were smoothed with a Gaussian filter of length 1.96 s (15 frames). The public dataset is distributed in MATLAB format and includes the processed activity matrices, state vectors, ROI centroid coordinates, cortical region labels, frame annotations, metadata, and an optional activity-based ROI inclusion flag (nonzero_ROI) used in the associated study.

### Data Records

#### Repository and overall organization

This dataset contains both processed and raw data recording sessions used in our previous study and is publicly available at the RIKEN NeuroData repository under accession ID 20260708-001 (DOI: 10.60178/cbs.20260708-001). It comprises processed wide-field two-photon calcium imaging data from the mouse cortex recorded across wakefulness, sleep (including NREM and REM sleep), and isoflurane anesthesia, together with spatial annotations, behavioral state labels, frame annotations, and metadata. The processed dataset contains the intermediate representations required for most downstream analyses, including fluorescence signals, deconvolved spike estimates, smoothed activity traces, and aligned annotations.

In addition, the repository provides the corresponding raw imaging movies in TIFF format and raw EEG/EMG recordings in MATLAB (.mat) format.

#### Recording sessions

The dataset contains 10 recording sessions in total: 6 wakefulness-sleep sessions and 4 wakefulness-anesthesia sessions. Each file corresponds to one continuous recording session. In the wakefulness-sleep sessions, activity was recorded across wakefulness and natural sleep, with post hoc classification into wakefulness, quiet wakefulness, NREM sleep, and REM sleep. In the wakefulness-anesthesia sessions, activity was recorded during wakefulness followed by isoflurane anesthesia. In the associated study^12^, each column in Tables S1 and S2 corresponds to a single session.

### File format

All shared data are provided in MATLAB (.mat) format. Each file contains processed neuronal activity data and associated annotations for one recording session. The data structure and variable definitions are described in the accompanying repository README.

#### Neuronal activity data

Processed neuronal activity is provided as matrices of size N × T (neurons × time points). For each recording session, the dataset includes three complementary activity representations: Δ F/F fluorescence signals (dFF), deconvolved spike estimates (spike_deconv), and Gaussian-smoothed deconvolved signals (spike_smoothed). These formats support a range of downstream analyses, from signal-level characterization to event-based and network-based analyses.

#### Behavioral state annotations

A behavioral state vector (state) is provided for each session and is temporally aligned with the neuronal activity matrices. In sleep recordings, the state annotations include wakefulness, quiet wakefulness, NREM sleep, and REM sleep. In anesthesia recordings, the state annotations include wakefulness and isoflurane anesthesia. State labels were assigned from simultaneously recorded EEG and EMG signals using a 4-s sliding window and post hoc classification.

### Spatial information

For each neuron, the dataset includes spatial annotations consisting of cortical region labels (ROIs.atlas) and centroid coordinates in imaging space (ROIs.Centroid). These variables enable analyses of neuronal population activity across spatially contiguous cortical areas within the same field of view.

### Frame annotations

Frame-level annotations are included to describe temporal segmentation and analysis windows. These variables include discontinuity indices (frame.boundary_ind) and state-specific frame indices (frame.used_frame). They allow users to identify valid continuous periods and reconstruct the subsets of frames used for state-specific analyses.

### Metadata

Additional session-level and animal-level metadata are provided in data_info and animal_info. These variables include recording condition, session type, and other information required for interpretation and reuse of the dataset.

#### ROI counts and quality-controlled populations

The number of detected and retained ROIs differs across sessions because the shared dataset preserves the outcome of session-specific preprocessing and quality control. In the wakefulness-sleep sessions, the number of originally detected ROIs ranged from 10,113 to 19,635, and the number retained after exclusion of contaminated ROIs ranged from 6,574 to 10,197. In the wakefulness-anesthesia sessions, the number of originally detected ROIs ranged from 7,571 to 10,400, and the number retained after quality control ranged from 4,337 to 7,350. These quality-controlled populations constitute the main shared dataset.

#### Analysis-specific variables

To facilitate direct reproduction of the network analyses reported in the associated study^12^, an optional activity-based neuron inclusion flag (nonzero_ROI) is provided. This flag identifies neurons that satisfied the additional activity criterion used in the published network analyses. Specifically, neurons were required to exhibit at least one Ca²⁺ event within every analyzed time window, where the analyzed windows consisted of 1,500 consecutive imaging frames (196s) for sleep recordings and 2,900 consecutive imaging frames (379s) for anesthesia recordings. Neurons that were silent in one or more analyzed windows were therefore excluded from the published network analyses. This variable is provided solely to facilitate reproduction of the published analyses; users performing independent analyses may instead use the full set of quality-controlled ROIs.

### Reuse notes

The shared dataset is intended for reuse in studies of cortical population dynamics across brain states, benchmarking of analytical pipelines, and comparative analyses of spatially distributed neuronal activity across physiological and pharmacological conditions. Because the dataset includes aligned activity traces, state annotations, and spatial coordinates, it is suitable for analyses spanning single-neuron, population, and spatially structured representations.

Variables included in each MATLAB (.mat) file

The variables included in each MATLAB (.mat) file are summarized in Table 1. Each file corresponds to one continuous recording session and contains processed neuronal activity data together with aligned annotations and metadata.

**Table 1.** Variables included in each MATLAB (.mat) file N, number of neurons (quality-controlled ROIs) in a session; T, number of imaging frames in that session. Each file contains one recording session from either the wakefulness-sleep dataset or the wakefulness-anesthesia dataset.

| Variable | Type / dimensions | Description | Notes |
| --- | --- | --- | --- |
| dFF | numeric matrix,<br>$\mathbf{N} \times \mathbf{T}$ | Neuropil-corrected<br>$\Delta F/F$ fluorescence<br>signals for all<br>quality-controlled<br>ROIs in a session | $\mathbf{N}$, number of neurons;<br>$\mathbf{T}$, number of imaging<br>frames |
| spike_deconv | numeric matrix,<br>$\mathbf{N} \times \mathbf{T}$ | Deconvolved<br>activity estimates<br>derived from dFF<br>using OASIS | Intended for event-based<br>and network analyses |
| spike_smoothed | numeric matrix,<br>$\mathbf{N} \times \mathbf{T}$ | Gaussian-smoothed<br>deconvolved activity<br>traces | Smoothing window: 15<br>frames (1.96 s at 7.65<br>Hz) |
| state | numeric vector, | Behavioral state | Sleep sessions include |
|  | length **T** | annotation aligned to imaging frames | wakefulness, quiet wakefulness, NREM sleep, and REM sleep; anesthesia sessions include wakefulness and isoflurane anesthesia |
| ROIs.atlas | string array, length **N** | Cortical region label assigned to each ROI | Region labels are based on cortical spatial annotation of the imaging field |
| ROIs.Centroid | numeric matrix, **N × 2** | Centroid coordinates of each ROI in imaging space | Used for spatial analyses across the field of view |
| frame.boundary_ind | numeric vector | Indices marking discontinuities or boundaries in the frame sequence | Useful for identifying valid continuous segments |
| frame.used_frame | cell array of numeric vectors (1 × n, double) representing frame indices | State-specific frame indices used for downstream analyses | Supports reconstruction of state-specific analysis windows |
| data_info.recording | string | Session-level metadata | Indicates the recording condition ("sleep" or "anesthesia") |
| data_info.circadian_phase | string | Session-level metadata | Indicates the recording condition ("Light" or "Dark") |
| animal_info | struct | Animal-level metadata | Includes animal information (age in weeks and sex) |
| nonzero_ROI | numeric vector,<br>$N \times 1$, logical | Binary flag indicating whether each ROI was included in the published network analyses. | Included only if at least one $\text{Ca}^{2+}$ event was detected in every analyzed window (1,500 frames for sleep; 2,900 frames for anesthesia). |

**Table 2.**
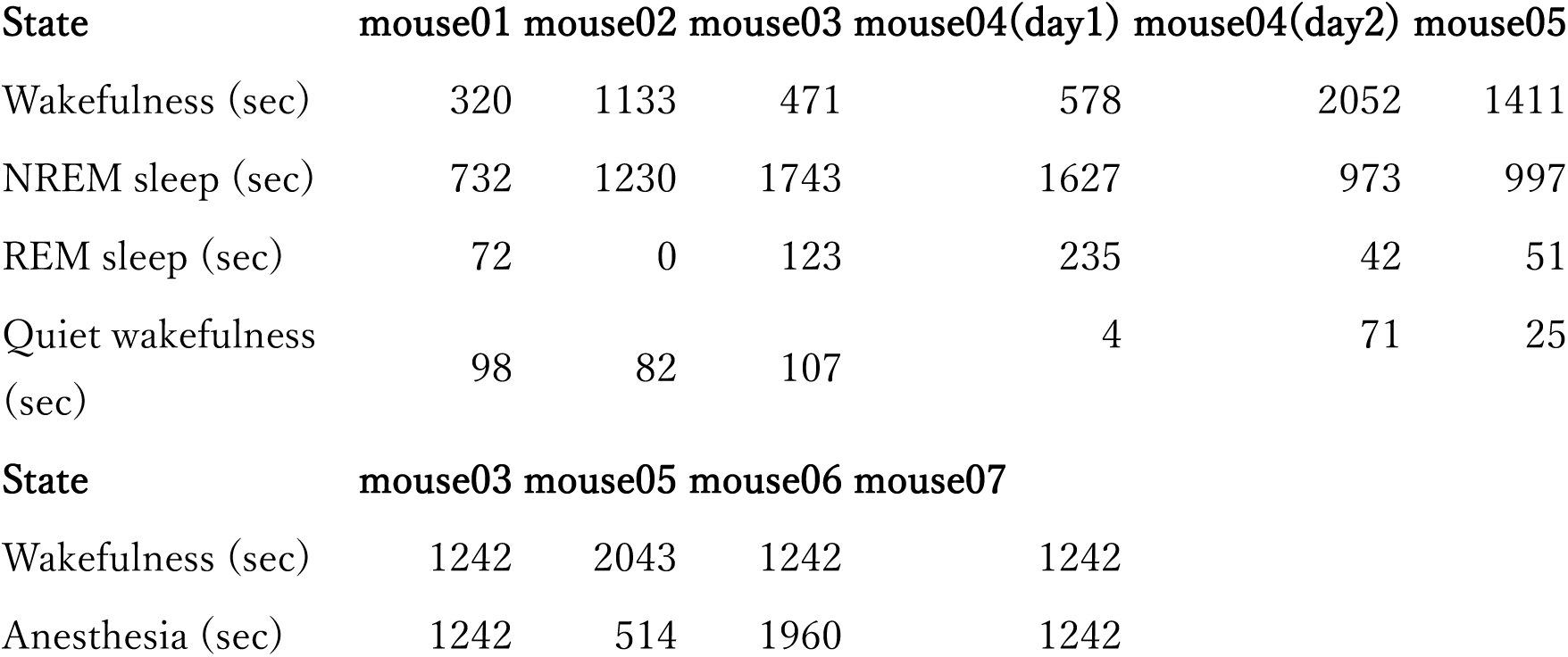
Cumulative duration of each behavioral state for each recording session in the sleep and anesthesia datasets.

Durations are reported in seconds.

Technical Validation

Data quality was assessed through standardized preprocessing, signal extraction, and quality-control procedures that were applied consistently across all recording sessions. Raw imaging movies were first corrected for motion using NoRMCorre^20^ and regions of interest (ROIs) corresponding to neuronal somata were identified using the low-computational-cost cell detection (LCCD) algorithm^21^. Neuropil-corrected fluorescence signals were then computed for each ROI, and Δ F/F traces were obtained using a sliding baseline defined as the 8th percentile within a ±30 s window, followed by additional baseline correction using the estimate_baseline function in Suite2P^26^. These procedures reduced motion-related and slow baseline artifacts and yielded stable fluorescence time series suitable for downstream analysis.

To exclude low-quality or contaminated ROIs, we applied multiple quality-control criteria reflecting both signal reliability and biological plausibility. For each ROI, the signal-to-noise ratio (SNR) of the Δ F/F trace was calculated in MATLAB, with signal defined as the component below 0.5 Hz and noise as the component above 0.5 Hz. ROIs were retained only if they satisfied all of the following conditions: (1) SNR within the top 90% of ROIs, (2) absence of obscuration by nearby blood vessels, (3) Δ F/F range between −1 and 10, (4) at least one calcium transient during recording, (5) neuropil fluorescence lower than somatic fluorescence, (6) fewer than 45% of frames with Δ F/F values below the mean, and (7) correlation between Δ F/F and the square root of the EMG signal below 0.2. Blood vessels were detected semi-automatically in ImageJ. ROIs that passed all criteria were retained in the shared dataset.

To provide additional activity representations for reuse, Δ F/F traces were deconvolved using OASIS^28^ and subsequently smoothed with a Gaussian filter of length 1.96 s (15 frames).The shared dataset therefore includes three complementary signal representations— Δ F/F fluorescence traces, deconvolved spike estimates, and Gaussian-smoothed deconvolved traces—allowing users to select the representation best suited to their analytical purpose. Representative examples of the field of view, ROI detection, raw Δ F/F traces, the effects of deconvolution and smoothing, and representative EEG/EMG recordings are shown in Fig. 1a–d. These panels illustrate the quality of segmentation and the consistency of signal processing across the analysis pipeline.

We further evaluated the consistency of signal quality across sessions and cortical regions using the distributions of SNR and standardized shot noise levels. As shown in Fig. 1e–h, these distributions were broadly consistent across recordings, indicating stable data quality across animals and experimental conditions. A subset of anesthesia recordings showed moderately elevated noise levels, but these differences did not substantially alter the overall distributions or compromise the usability of the dataset, because all recordings were processed and filtered using the same quality-control criteria. Region-wise analyses likewise showed comparable signal quality across cortical areas.

Because some downstream analyses require the same set of active neurons across multiple time windows, we additionally provide an optional activity-based inclusion flag (nonzero_ROI) that identifies neurons exhibiting at least one Ca²⁺ event in all analyzed windows of a session. This variable facilitates exact reproduction of the published network analyses, while the main dataset retains the full quality-controlled ROI population for broader reuse. Taken together, these preprocessing, filtering, and validation steps indicate that the dataset provides reliable measurements of large-scale cortical population activity suitable for quantitative analysis, benchmarking, and method development across brain states.

Usage Notes

This dataset is organized as session-based data files, with one file corresponding to one continuous recording session. The shared data are suitable for a broad range of downstream analyses, including characterization of state-dependent population activity, spatial analyses across cortical areas, benchmarking of preprocessing and inference methods, and reproduction or extension of the network analyses reported in the associated study. The repository includes both processed MATLAB files, corresponding raw imaging movies (TIFF), and raw electrophysiological recordings (EEG/EMG) in MATLAB format, allowing users to perform analyses at either the processed-signal level or the raw image level, including alternative motion correction, ROI segmentation, and fluorescence extraction. Representative activity movies are also available through the Additional Notes section of the repository, allowing users to visually inspect examples of large-scale cortical population dynamics.

Users can select among three activity representations according to their analytical goals. The dFF variable is most appropriate for analyses focused on fluorescence dynamics and signal-level properties, whereas spike_deconv is better suited for event-based and network analyses that require temporally sharpened activity estimates. The spike_smoothed variable may be useful when more robust population-level summaries are desired. Behavioral state annotations (state) are aligned to imaging frames and can be used directly for state-specific analyses. In sleep sessions, the annotations distinguish wakefulness, quiet wakefulness, NREM sleep, and REM sleep; in anesthesia sessions, the annotations distinguish wakefulness and isoflurane anesthesia.

For most reuse scenarios, we recommend starting from the full quality-controlled ROI population. An additional variable, nonzero_ROI, is provided for users who wish to reproduce the published network analyses exactly. This variable identifies neurons that exhibited at least one detected Ca²⁺ event in every analyzed window of a session and was introduced to keep the node set constant within each session during network comparisons. Because this activity-based filtering is more restrictive than the main quality-control procedure, analyses using nonzero_ROI should be interpreted as being based on an activity-selected subset of neurons rather than on the full recorded population.

Users should also note several dataset-specific considerations when comparing brain states. First, wakefulness-sleep sessions and wakefulness-anesthesia sessions were acquired as separate session types, so direct comparisons between sleep and anesthesia should be made with caution. Second, in sleep sessions, REM sleep and quiet wakefulness were annotated but were relatively rare, and the associated study therefore focused primarily on wakefulness and NREM sleep for network analyses. Users interested in rare-state analyses should consider the limited duration of these epochs on a session-by-session basis.

The dataset includes spatial coordinates and cortical region labels for each ROI, enabling analyses at multiple levels of spatial organization, from single-neuron activity to cross-area population structure. Cortical region labels (ROIs.atlas) were assigned using approximate atlas registration and should therefore be interpreted as reference information, particularly for neurons located near cortical area boundaries. The recordings were obtained at 7.65 Hz from layer 2/3 neurons within a contiguous wide field of view, making the dataset particularly well suited for analyses of mesoscale spatial organization built directly from cellular-resolution activity. However, users should keep in mind that the dataset is restricted to recordings from layer 2/3 cortical populations and does not include simultaneous sampling across cortical depths.

Additional information on the dataset structure, variable definitions, file organization, and example analysis workflows is provided in the accompanying README documentation. The README also includes example code for loading the dataset in MATLAB and Python, together with a typical workflow illustrating how to identify behavioral states and perform downstream analyses.

## Code Availability

Code used to reproduce the network analyses reported in the associated study is available at GitHub (https://github.com/oizumi-lab/mouse_network_2P). No custom code is required to access the dataset described in this Data Descriptor. The shared data files can be loaded and analyzed using standard MATLAB workflows.

## Data Availability

The dataset described in this study is publicly available at the RIKEN NeuroData repository under accession ID 20260708-001 (DOI: 10.60178/cbs.20260708-001). The repository contains both processed and raw data, including wide-field two-photon calcium imaging data, raw TIFF image sequences, raw EEG/EMG recordings in MATLAB (.mat) format, behavioral state annotations, spatial annotations, and metadata for all recording sessions included in this study. The processed dataset includes Δ F/F fluorescence signals, deconvolved spike estimates, Gaussian-smoothed activity traces.

## Acknowledgements

We thank Yoshihito Saito and Yusuke Atsumi for valuable comments and discussions. This research was supported by Japan Society for the Promotion of Science KAKENHI grants JP19J01973 and JP24K18246 (to I.O.), JP20H05775 and JP24H02313 (to M.M.), JP20H05712 and JP23H04834 (to M.O.), and JP24KJ0762 (to D.K.); AMED-Brain/Minds 1.0 project JP15dm0207001 and AMED-Brain/Minds 2.0 project JP23wm0625001 (to M.M.); KAO Corp. (to M.M.); the Toray Science Foundation (to M.M.); Japan Science and Technology Agency CREST grant JPMJCR1864 (to M.O.); JST SPRING grant JPMJSP2108 (to D.K.); JST Moonshot R&D grant JPMJMS2012 (to M.O.); RIKEN Incentive Research Projects (to M.M.); and the RIKEN Special Postdoctoral Researcher Program (to I.O.).

## Author Contributions

Conceptualization: Oomoto, Murayama Data curation: Oomoto, Kiyooka Formal analysis: Oomoto, Kiyooka Investigation: Oomoto, Kiyooka Methodology: Oomoto, Murayama Supervision: Oizumi, Murayama

Writing – original draft: Oomoto Writing – review & editing: all authors

## References

1. Harris, K. D. & Thiele, A. Cortical state and attention. Nat. Rev. Neurosci. 12, 509–523 (2011).

2. Brown, E. N., Lydic, R. & Schiff, N. D. General Anesthesia, Sleep, and Coma. N. Engl. J. Med. 363, 2638–2650 (2010).

3. Vyazovskiy, V. V. et al. Cortical Firing and Sleep Homeostasis. Neuron 63, 865–878 (2009).

4. Issa, E. B. & Wang, X. Sensory responses during sleep in primate primary and secondary auditory cortex. J. Neurosci. Off. J. Soc. Neurosci. 28, 14467–14480 (2008).

5. Nir, Y., Vyazovskiy, V. V., Cirelli, C., Banks, M. I. & Tononi, G. Auditory Responses and Stimulus-Specific Adaptation in Rat Auditory Cortex are Preserved Across NREM and REM Sleep. Cereb. Cortex 25, 1362–1378 (2015).

6. Krom, A. J. et al. Anesthesia-induced loss of consciousness disrupts auditory responses beyond primary cortex. Proc. Natl. Acad. Sci. 117, 11770–11780 (2020).

7. Steriade, M., McCormick, D. A. & Sejnowski, T. J. Thalamocortical Oscillations in the Sleeping and Aroused Brain. Science 262, 679–685 (1993).

8. Steriade, M., Nunez, A. & Amzica, F. A novel slow (< 1 Hz) oscillation of neocortical neurons in vivo: depolarizing and hyperpolarizing components. J. Neurosci. 13, 3252– 3265 (1993).

9. Vyazovskiy, V. V. et al. Local sleep in awake rats. Nature 472, 443–447 (2011).

10. Lewis, L. D., et al. Rapid fragmentation of neuronal networks at the onset of propofol-induced unconsciousness. Proc. Natl. Acad. Sci. 109, E3377–E3386 (2012).

11. Ota, K. et al. Fast, cell-resolution, contiguous-wide two-photon imaging to reveal functional network architectures across multi-modal cortical areas. Neuron 109, 1810–1824.e9 (2021).

12. Kiyooka, D. et al. Single-cell resolution functional networks during unconsciousness are segregated into spatially intermixed modules. Cell Rep. 45, 116902 (2026).

13. Saito, Y., Osako, Y. & Murayama, M. Unraveling the neural code: analysis of large-scale two-photon microscopy data. Microscopy 74, 146–163 (2025).

14. Ota, K., Uwamori, H., Ode, T. & Murayama, M. Breaking trade-offs: Development of fast, high-resolution, wide-field two-photon microscopes to reveal the computational principles of the brain. Neurosci. Res. 179, 3–14 (2022).

15. Shiba, Y. et al. Allogeneic transplantation of iPS cell-derived cardiomyocytes regenerates primate hearts. Nature 538, 388–391 (2016).

16. Oomoto, I. et al. Protocol for cortical-wide field-of-view two-photon imaging with quick neonatal adeno-associated virus injection. STAR Protoc. 2, 101007 (2021).

17. Kawai, S., Takagi, Y., Kaneko, S. & Kurosawa, T. Effect of Three Types of Mixed Anesthetic Agents Alternate to Ketamine in Mice. Exp. Anim. 60, 481–487 (2011).

18. Miyamoto, D. et al. Top-down cortical input during NREM sleep consolidates perceptual memory. Science 352, 1315–1318 (2016).

19. Saito, Y. et al. Amygdalo-cortical dialogue underlies memory enhancement by emotional association. Neuron 113, 931–948.e7 (2025).

20. Pnevmatikakis, E. A. & Giovannucci, A. NoRMCorre: An online algorithm for piecewise rigid motion correction of calcium imaging data. J. Neurosci. Methods 291, 83–94 (2017).

21. Ito, T. et al. Low computational-cost cell detection method for calcium imaging data. Neurosci. Res. 179, 39–50 (2022).

22. Kerlin, A. M., Andermann, M. L., Berezovskii, V. K. & Reid, R. C. Broadly Tuned Response Properties of Diverse Inhibitory Neuron Subtypes in Mouse Visual Cortex. Neuron 67, 858–871 (2010).

23. Peron, S. et al. Recurrent interactions in local cortical circuits. Nature 579, 256–259 (2020).

24. Chen, T.-W. et al. Ultrasensitive fluorescent proteins for imaging neuronal activity. Nature 499, 295–300 (2013).

25. Dombeck, D. A., Khabbaz, A. N., Collman, F., Adelman, T. L. & Tank, D. W. Imaging Large-Scale Neural Activity with Cellular Resolution in Awake, Mobile Mice. Neuron 56, 43–57 (2007).

26. Pachitariu, M. et al. Suite2p: beyond 10,000 neurons with standard two-photon microscopy. 061507 Preprint at 10.1101/061507 (2017).

27. Rupprecht, P. et al. A database and deep learning toolbox for noise-optimized, generalized spike inference from calcium imaging. Nat. Neurosci. 24, 1324–1337 (2021).

28. Friedrich, J., Zhou, P. & Paninski, L. Fast online deconvolution of calcium imaging data. PLOS Comput. Biol. 13, e1005423 (2017).

